# Hippocampal microstructure, but not macrostructure, mediates age differences in episodic memory

**DOI:** 10.1101/2022.07.13.499977

**Authors:** Kirolos Ibrahim, Ilana J. Bennett

**Author notes:** Author Notes. Data Availability Statement: The data that support the findings of this study are available from the corresponding author upon request. Declarations of interest: None. Corresponding author: Ilana J. Bennett Department of Psychology University of California, Riverside 900 University Avenue Riverside, CA 92521.

## Abstract

Separate unimodal magnetic resonance imaging (MRI) literatures have shown that hippocampal gray matter macrostructure (volume) and microstructure (diffusion) decline in aging and relate to episodic memory performance, with multimodal MRI studies reporting that episodic memory may be better explained by a combination of these metrics. However, these effects are often assessed independent of age or only within older adults and therefore do not address whether these distinct modalities mediate the effect of age on episodic memory. Here, we simultaneously examined the unique and joint contribution of hippocampal volume and diffusion to age-related differences in episodic memory in 83 younger and 61 older adults who underwent a T1-and diffusion-weighted MRI and completed the Rey Auditory Verbal Learning Test. As expected, older age was significantly related to smaller volume and higher diffusion (intracellular, dispersion, and free) in bilateral hippocampus and to worse episodic memory performance (immediate and delayed free recall, recognition). Structural equation modelling revealed that the age-memory relationship was significantly mediated by hippocampal diffusion, but not volume. A non-significant influential indirect effect further revealed that the structural metrics did not jointly mediate the age-memory relationship. Together, these findings indicate that hippocampal microstructure uniquely contributes to age-related differences in episodic memory and suggest that volume and diffusion capture distinct neurobiological properties of hippocampal gray matter.

## Introduction

The neurocognitive aging field seeks to identify the neural basis of age-related declines in various cognitive abilities. A key function known to decline in otherwise cognitively normal older adults is the ability to remember past events (Craik, 1994; Nyberg, Backman, et al., 1996). These episodic memories are often assessed using list learning tasks, such as the Rey Auditory Verbal Learning Task (RAVLT)(Rey, 1941), in which participants study a list of words and are later tested on their ability to generate those words without cues (free recall) or identify them among distractor words (recognition). Older adults consistently recall fewer words than younger adults, particularly after a delay, but there are mixed findings regarding age effects for recognition memory (Craik & McDowd, 1987; Parker et al., 2004; Rhodes et al., 2019). These age-related differences in episodic memory are often attributed to degradation of the hippocampus in older adults (Leal & Yassa, 2015; van Petten, 2004), which can be assessed using structural magnetic resonance imaging (MRI). Whereas most studies report on a single MRI modality, multimodal approaches are needed to determine which modalities are more sensitive to the effect of age on memory performance.

One of the most commonly used structural MRI modalities is T1-weighted imaging, which measures differences in tissue contrast and can be used to assess macrostructural properties of individual brain regions, such as their size or shape (Orrison et al., 1995). Unimodal T1-weighted MRI studies have consistently shown age-related decreases in volume of the hippocampus (Du et al., 2006; Nobis et al., 2019; Raz et al., 2005; Walhovd et al., 2005) and that smaller hippocampal volume relates to worse delayed free recall (Bruno et al., 2016; de Leon et al., 1997; Golomb et al., 1994; Lye et al., 2004) and recognition memory (Bender et al., 2013; Bennett et al., 2019; Shing et al., 2011). However, these volume-memory relationships are primarily seen within older adults whereas studies of adults across the lifespan do not observe significant relationships between hippocampal volume and episodic memory (Raz et al., 1998; Rodrigue & Raz, 2004; Sullivan et al., 1995; Tisserand et al., 2000).

A complementary structural MRI modality is diffusion-weighted imaging, which measures the movement of molecular water and can be used to assess microstructural properties of tissue, such as the presence and organization of neurons and glia (Afzali et al., 2021; Beaulieu, 2002). Neurite Orientation Dispersion and Density Imaging (NODDI) is a multi-compartment modelling approach that yields separate estimates of diffusion within (intracellular) and between (dispersion) cells and from non-cellular sources (free)(Zhang et al., 2012). These NODDI metrics are more sensitive to age differences in diffusion in gray matter than single tensor metrics (Venkatesh et al., 2020). NODDI studies have shown age-related increases in all diffusion metrics in the hippocampus (Franco et al., 2021; Metzler-Baddeley et al., 2019; Nazeri et al., 2015; Radhakrishnan et al., 2020; Venkatesh et al., 2020, 2021) and that higher hippocampal diffusion relates to worse delayed free recall in younger and older adults (Radhakrishnan et al., 2020; Venkatesh et al., 2021). Thus, whereas hippocampal macrostructure (volume) only relates to episodic memory performance within older adults, hippocampal microstructure (diffusion) is sensitive to individual differences in memory performance across the adult lifespan. What remains untested is the extent to which either of these MRI modalities in hippocampal gray matter mediate the effect of age on episodic memory.

An important follow up question is whether macrostructural and microstructural properties of hippocampal gray matter are differentially sensitive to effects of age on episodic memory. The multimodal MRI literature has primarily provided evidence that diffusion captures unique variance that is more sensitive to episodic memory than volume, although these studies have been limited to older adults or assessed the structure-memory relationships independent of age. For example, one study that compared these MRI modalities in the same group of older adults found that single tensor metrics of hippocampal diffusion, but not hippocampal volume, predicted memory performance (den Heijer et al., 2012). Another study reported that episodic memory performance across younger and older adults was better predicted when multi-compartment diffusion in the hippocampus was added to a model of hippocampal volume and single tensor diffusion metrics (Radhakrishnan et al., 2022). A third multimodal MRI study instead focused on their shared variance, finding that episodic memory in older adults was related to a latent construct of structural MRI modalities, including T1– and diffusion-weighted metrics, in the hippocampus (Köhncke et al., 2021). The combination of unique and shared variance in memory performance may indicate that volume and diffusion capture at least some different underlying neurobiological substrate(s), such as neurodegeneration, demyelination, or inflammation, or perhaps the same substrates but to different degrees.

Addressing the gaps of prior work, the current study examined the extent to which hippocampal macrostructure (volume) and microstructure (diffusion) act independently and together to mediate the effect of age on multiple forms of episodic memory (immediate free recall, delayed free recall, yes/no recognition) in a large sample of younger and older adults (n = 144). These complex relationships were tested using structural equation modeling (SEM), which is a powerful tool that allows for simultaneous examination of unique (specific indirect effects) and joint (influential indirect effect) mediational influences of multiple MRI metrics. Specific indirect effects were expected for diffusion, with weaker or non-significant effects for volume. A significant influential indirect effect would further indicate that these distinct MRI modalities explain shared variance in the effect of age on memory performance, with a “causal” pathway linking microstructure to macrostructure (or vice versa), as would be expected if they capture common neurobiological substrates. Additional analyses used raw volume instead of normalized volume as normalization methods have been shown to influence volume-memory relationships in older adults (van Petten, 2004) and separate models for each memory metric as prior work was largely limited to free recall. An exploratory analysis reran the original model within older adults only for comparison to prior work.

## Methods

### Participants

Eighty-nine healthy younger adults and 83 older adults were recruited from the University of California, Riverside undergraduate research pool and surrounding Riverside community, respectively. All participants were screened for conditions that would prevent them from safely entering an MRI scanner (e.g., ferrous mental implants, claustrophobia, pregnancy). Participants were excluded for having missing data (1 younger, 6 older) data, poor general cognition assessed using the Mini-Mental State Exam (scores <26; 3 younger)(Folstein et al., 1975) or Montreal Cognitive Assessment (scores <21; 2 older)(Pendlebury et al., 2013), significant lesions in bilateral hippocampus (6 older), uncorrectable MRI registration or segmentation errors (1 younger, 3 older), MRI artifacts (values exceeding 3 standard deviations from the sample mean; 2 older), or poor episodic memory performance (RAVLT recognition >3 standard deviations below the sample mean; 1 younger, 3 older). The final sample included 83 younger (18-29 years) and 61 older (65-86 years) adults (see Table 1). All participants gave consent and received either course credit or monetary compensation for their participation.

**Table 1.**
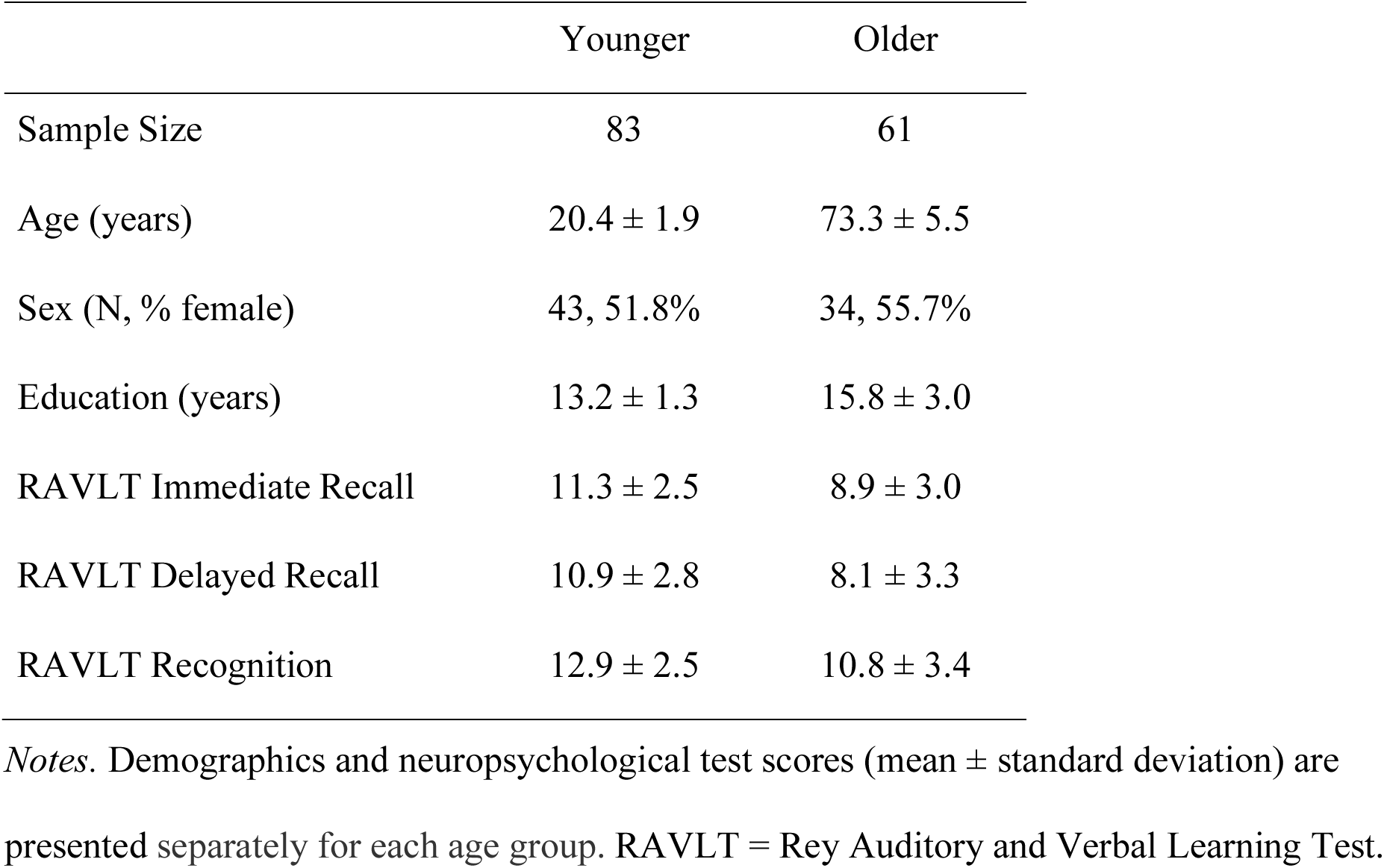
Demographic and neuropsychological test data.

### Episodic Memory Task

Participants completed the Rey Auditory Verbal Learning Task (RAVLT)(Rey, 1941) to assess free recall and yes/no recognition. On each on five recall trials, the experimenter read the same list of 15 common words (List A) after which the participant was asked to freely recall as many words as possible. On a subsequent interference trial, the experimenter read a different list of 15 common words (List B) that the participant had to freely recall. Immediate and delayed recall scores were calculated as the number of words correctly recalled from List A immediately after the interference trial and after a 20-minute delay, respectively. The experimenter then read a list of 50 words and the participant had to indicate if they were (“yes”) or were not (“no”) from List A. A recognition score was calculated as the difference between the number of correctly (hits) and incorrectly (false alarms) identified List A words.

### Imaging Acquisition Protocol

Imaging data were acquired using a 3-T Siemens MRI (Siemens Healthineers, Malvern, PA) scanner with a 32-channel receive-only head coil at the Center for Advanced Neuroimaging at the University of California, Riverside. Head movement was minimized by placing a fitted padding around each participant#s head.

A single T1-weighted high-resolution magnetization prepared rapid acquisition gradient echo (MP-RAGE) sequence was acquired using the following parameters: echo time (TE)/repetition time (TR) = 2.72/2400 ms, 208 axial slices, field of view = 256 × 256 × 208 mm, flip angle = 8 degrees, and spatial resolution = 0.8 mm^3^.

A pair of diffusion-weighted scans with opposite phase encoding directions were also acquired. Each scan had two gradient strengths (b = 1500 and 3000 s/mm^2^) applied in 64 orthogonal directions, with six images having no diffusion weighting (b = 0) and using the following acquisition parameters: TE/TR = 102/3500 ms, field of view = 212 × 182 mm, 72 interleaved slices, and spatial resolution = 1.7 mm^3^.

### Region of Interest Segmentation

Bilateral hippocampus was automatically segmented on each participant#s MP-RAGE using FMRIB Software Library (FSL)(Jenkinson et al., 2012) Integrated Registration and Segmentation Tool (FIRST)(Patenaude et al., 2011). The three-stage affine registration flag was used to optimize fitting of the standard space model to subject space. Outputs were visually inspected by trained researchers, and participants were excluded for excessive (>1 voxel) and uncorrectable over– or under-fitting.

### Volumetric Data Processing

For each participant, raw hippocampal volumes were calculated separately in each hemisphere from the FIRST-segmented structures (Volumeraw). Individual differences in brain size were then corrected using the residual normalization method (Jack et al., 1989). Intracranial volume was measured for each participant using the Estimated Total Intracranial Volume (eTIVindiv) generated by FreeSurfer (v.6.0; http://surfer.nmr.mgh.harvard.edu) and then averaged within younger participants (eTIVmean). The effect of brain size on hippocampal volume was estimated using the slope of the regression line between eTIVindiv and Volumeraw, separately for each hemisphere (*β*) within younger participants. Normalized volumes (Volumenorm) were then calculated separately for bilateral hippocampus in each participant using the equation: Volumenorm = Volumeraw − β (eTIVindiv − eTIVmean).

### Diffusion Data Processing

For each participant, diffusion data were pre-processed using Analysis of Functional Neuro Images (AFNI)(Cox & Hyde, 1997) to remove non-brain tissue and generate a whole brain mask. FSL#s Topup was then used to generate a field map and Eddy was used to correct for motion, eddy-current induced distortions, and susceptibility induced distortions.

Pre-processed diffusion data were analyzed using the NODDI MATLAB toolbox (http://nitrc.org/projects/noddi_toolbox), which generates voxel-wise estimates of three metrics thought to reflect different sources of the diffusion signal (Tariq et al., 2016; Zhang et al., 2012). For better modeling within gray matter, the intrinsic diffusivity metric was adjusted to 1.1 μM^3^ (Guerrero et al., 2019). Intracellular diffusion (also known as intracellular volume fraction, fICVF) is modelled as a set of sticks, dispersion of diffusion (also known as orientation dispersion index, ODI) is modelled as the dispersion of those sticks, and free diffusion (also known as fraction of isotropic diffusion, fISO) is modeled as an isotropic sphere.

These diffusion metrics were extracted from bilateral hippocampus for each participant. Segmented hippocampus masks were registered to diffusion space by applying the inverse of a rigid body transformation (6 degrees of freedom, boundary-based registration cost function) from a linear alignment between the brain extracted diffusion (distortion-corrected average b0) and MP-RAGE images. Mean diffusion metrics were then calculated separately for each hemisphere by multiplying each registered hippocampus mask by the corresponding voxel-wise NODDI image and taking the average across voxels. All diffusion metrics were limited to voxels with sufficient signal (excluding voxels with intracellular diffusion >0.99)(Emmenegger et al., 2021) and the intracellular and dispersion of diffusion metrics were further limited to voxels with sufficient cellular sources of the diffusion signal (excluding voxels with free diffusion >0.9).

## Data Analysis

### Latent factor construction

Latent constructs of hippocampal volume and diffusion and episodic memory were generated using AMOS by SPSS (version 27.0, IBM, 2020). The latent volume construct was built from measures of normalized hippocampal volume from each hemisphere. The latent diffusion construct was built from latent constructs of the hippocampal NODDI metrics (intracellular, dispersion, free), each of which was built from the corresponding mean diffusion metrics from each hemisphere. Finally, the latent construct of episodic memory was built from the RAVLT immediate and delayed free recall and recognition scores. A correlational latent variables model was used to determine whether the measured variables significantly represented their respective latent constructs.

### Structural equation modeling

To assess whether hippocampal volume and diffusion uniquely or jointly mediate the effect of age on episodic memory performance, we conducted structural equation models using data from all younger and older participants (or older adults only) with AMOS by SPSS (see Figure 1). The models depicted two specific indirect pathways (separately for latent constructs of diffusion and volume) as well as an influential indirect pathway (from the latent construct of diffusion to volume) between age and the latent memory construct (or each memory metric separately). The influential indirect pathway was calculated as the product of the direct effect estimates of age on diffusion, diffusion on volume, and volume on memory. Prior to model fitting and analyses, age was mean centered and all diffusion metrics were multiplied by –1 to match the direction of age effects across modalities. Modification indices were used to adjust the model if necessary.

**Figure 1.**
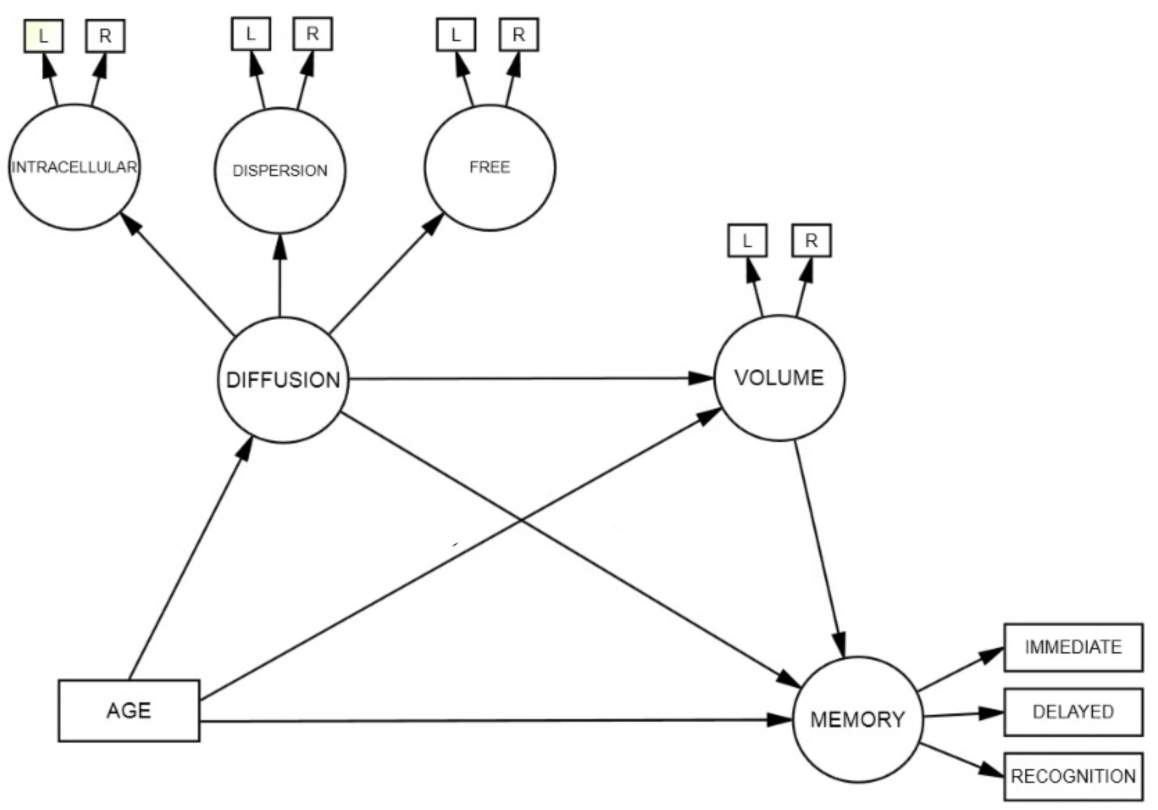
Structural equation model used to test the mediating effects of hippocampal diffusion (intracellular, dispersion, free) and volume on age-related differences in episodic memory performance (immediate free recall, delayed free recall, recognition) in younger and older participants. Circles represent latent variables and rectangles represent measured variables. Bolded values depict significant path estimates. L/R = left/right hemisphere.

Standardized path estimates were derived from bootstrapping using 5,000 iterations with 95% confidence intervals. Model fit was evaluated using established thresholds across several indices: non-significant chi-square (*X*^2^) value, root mean square error of approximation (RMSEA) ≤ 0.05, comparative fit index (CFI) > 0.90, and standardized root mean residual (SRMR) < 0.08 (Hu & Bentler, 1999; Raykov & Marcoulides, 2006). Improvements in model fit were assessed by comparing Akaike information criterion (AIC) and Bayesian information criterion (BIC) measures, with smaller values indicating better fit. Specific indirect effects were tested with the James and Brett (1984) method, by which a significant indirect effect is sufficient to support an indirect relationship by means of an intermediate variable.

## Results

### Effect of Age on Measured Variables

The effect of age on each measured variable was assessed using separate Pearson correlations (Table 2). Significant effects survived Bonferroni correction for 11 comparisons per measure, *p* < 0.0045. Results revealed that older age was significantly related to higher hippocampal diffusion, smaller hippocampal volume, and worse episodic memory performance. Moreover, all memory scores were significantly related to all hippocampal diffusion metrics, but not to hippocampal volume.

**Table 2.** Correlation coefficients for measured variables.

|  | Age | 1 | 2 | 3 | 4 | 5 | 6 | 7 | 8 | 9 | 10 |
| --- | --- | --- | --- | --- | --- | --- | --- | --- | --- | --- | --- |
| 1. Intracellular R | **-0.43** |  |  |  |  |  |  |  |  |  |  |
| 2. Intracellular L | **-0.51** | **0.88** |  |  |  |  |  |  |  |  |  |
| 3. Dispersion R | **-0.68** | **0.73** | **0.70** |  |  |  |  |  |  |  |  |
| 4. Dispersion L | **-0.66** | **0.65** | **0.75** | **0.86** |  |  |  |  |  |  |  |
| 5. Free R | **-0.37** | **0.86** | **0.76** | **0.66** | **0.61** |  |  |  |  |  |  |
| 6. Free L | **-0.47** | **0.78** | **0.88** | **0.65** | **0.71** | **0.81** |  |  |  |  |  |
| 7. Volume R | **-0.48** | 0.06 | 0.16 | **0.32** | **0.31** | 0.05 | 0.13 |  |  |  |  |
| 8. Volume L | **-0.41** | 0.08 | 0.08 | **0.31** | **0.34** | 0.14 | 0.10 | **0.69** |  |  |  |
| 9. Recognition | **-0.34** | **0.36** | **0.41** | **0.41** | **0.42** | **0.28** | **0.39** | 0.08 | 0.09 |  |  |
| 10. Immediate recall | **-0.40** | **0.40** | **0.46** | **0.40** | **0.41** | **0.33** | **0.41** | 0.13 | 0.07 | **0.60** |  |
| 11. Delayed recall | **-0.41** | **0.42** | **0.52** | **0.42** | **0.48** | **0.36** | **0.47** | 0.15 | 0.11 | **0.66** | **0.84** |

*Notes.* Correlation coefficients are shown for comparisons among age, hippocampal diffusion and volume from each hemisphere (L = left, R = right), and memory performance. Significant effects survived Bonferroni correction for 11 comparisons per measure (bolded; $p < 0.0045$ ).

### Latent Factor Analysis

Prior to model fitting, a correlational latent variables model confirmed that all measured variables significantly represented their respective latent constructs, as seen by significant standardized loading scores, *p*s < 0.01 (Table 3). As such, latent constructs for hippocampal diffusion, hippocampal volume, and episodic memory were used in the subsequent mediation analyses.

**Table 3.** Standardized measurement loading onto respective factors.

| Latent construct | Measured variable | Loading |
| --- | --- | --- |
|  | Age | 1.00 |
| Diffusion | Intracellular diffusion | 0.96 |
|  | Dispersion of diffusion | 0.82 |
|  | Free diffusion | 1.00 |
| Intracellular diffusion | Left hippocampus | 0.90 |
|  | Right hippocampus | 0.97 |
| Dispersion of diffusion | Left hippocampus | 0.91 |
|  | Right hippocampus | 0.94 |
| Free diffusion | Left hippocampus | 0.86 |
|  | Right hippocampus | 0.94 |
| Volume | Left hippocampus | 0.71 |
|  | Right hippocampus | 0.98 |
| Episodic memory | Delayed recall | 0.96 |
|  | Immediate recall | 0.87 |
|  | Recognition | 0.69 |

*Notes.* All loadings were significant at $p < 0.01$ .

## Structural Equation Models

### Original model in all participants

The original SEM in all participants (Figure 1) initially resulted in fits (AIC = 288.3, BIC = 383.3) that were improved after implementing a modification index correlating the residuals of right hemisphere hippocampal intracellular diffusion and right hemisphere hippocampal free diffusion (AIC = 207.0, BIC = 305.0). This resulted in acceptable model fits according to standard thresholds for some indices (CFI), but not others (*X*^2^, RMSEA, SRMR).

As seen in Table 4, significant path effects were seen for the total effect of age on memory, but not the direct effect of age on memory after accounting for the mediators. The specific indirect effect on the age-memory relationship was significant for hippocampal diffusion, but not normalized hippocampal volume. Moreover, the influential indirect pathway capturing the joint mediating effects of hippocampal diffusion and volume on the age-memory relationship was not significant.

**Table 4.**
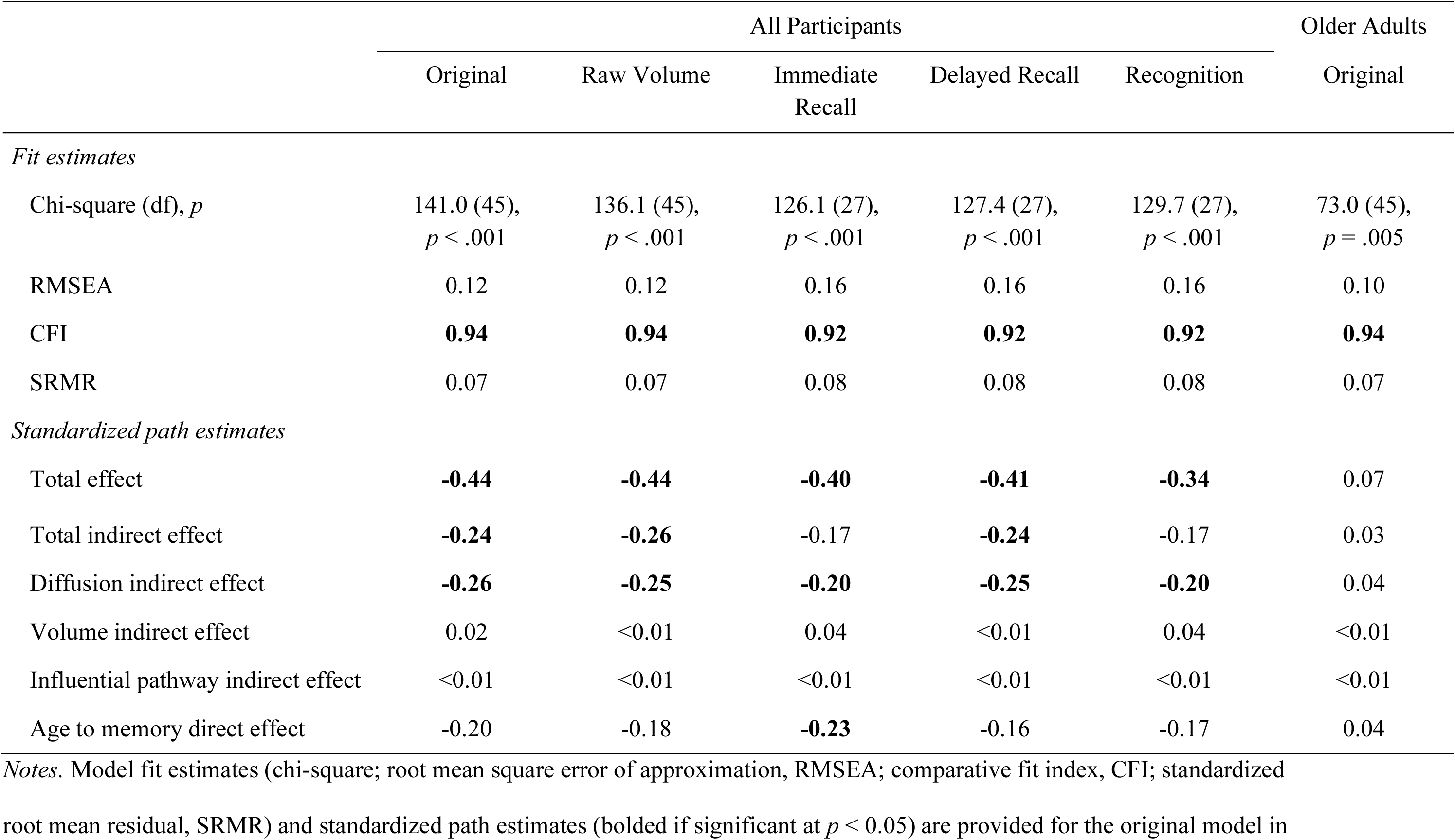
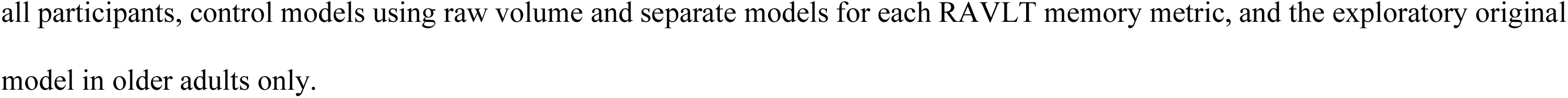
Structural equation model fit and standardized path estimates.

|  | All Participants |  |  |  |  | Older Adults |
| --- | --- | --- | --- | --- | --- | --- |
|  | Original | Raw Volume | Immediate Recall | Delayed Recall | Recognition | Original |
| *Fit estimates* |  |  |  |  |  |  |
| Chi-square (df), *p* | 141.0 (45),<br>*p* < .001 | 136.1 (45),<br>*p* < .001 | 126.1 (27),<br>*p* < .001 | 127.4 (27),<br>*p* < .001 | 129.7 (27),<br>*p* < .001 | 73.0 (45),<br>*p* = .005 |
| RMSEA | 0.12 | 0.12 | 0.16 | 0.16 | 0.16 | 0.10 |
| CFI | **0.94** | **0.94** | **0.92** | **0.92** | **0.92** | **0.94** |
| SRMR | 0.07 | 0.07 | 0.08 | 0.08 | 0.08 | 0.07 |
| *Standardized path estimates* |  |  |  |  |  |  |
| Total effect | **-0.44** | **-0.44** | **-0.40** | **-0.41** | **-0.34** | 0.07 |
| Total indirect effect | **-0.24** | **-0.26** | -0.17 | **-0.24** | -0.17 | 0.03 |
| Diffusion indirect effect | **-0.26** | **-0.25** | **-0.20** | **-0.25** | **-0.20** | 0.04 |
| Volume indirect effect | 0.02 | <0.01 | 0.04 | <0.01 | 0.04 | <0.01 |
| Influential pathway indirect effect | <0.01 | <0.01 | <0.01 | <0.01 | <0.01 | <0.01 |
| Age to memory direct effect | -0.20 | -0.18 | **-0.23** | -0.16 | -0.17 | 0.04 |

*Notes.* Model fit estimates (chi-square; root mean square error of approximation, RMSEA; comparative fit index, CFI; standardized root mean residual, SRMR) and standardized path estimates (bolded if significant at $p < 0.05$ ) are provided for the original model in
all participants, control models using raw volume and separate models for each RAVLT memory metric, and the exploratory original model in older adults only.

### Control model using raw volume

Results of the original model in all participants remained unchanged when using raw volume instead of normalized volume. Model fits were comparable to the original model (AIC = 202.1, BIC = 300.1) and the same standardized path effects were significant (Table 4).

### Control models for each memory metric

Results of the original model in all participants remained unchanged when separate models were conducted for each memory measure rather than using the latent memory construct. Model fits were comparable to the original model (immediate free recall: AIC = 182.1, BIC = 265.3; delayed free recall: AIC = 183.4, BIC = 266.5; recognition: AIC = 185.7, BIC = 268.9) and similar standardized path effects were significant, except that the total indirect effect was not significant for the immediate free recall or recognition models and the direct effect was significant for the immediate free recall model (Table 4).

### Exploratory model in older adults only

Given that relationships between hippocampal volume and episodic memory had been primarily reported in studies of older adults, rather than adults across the lifespan, the original SEM was run again using data from only older adults. Using the same modification index, model fits for this exploratory SEM were comparable to the original model (AIC = 139.0, BIC = 208.6), with acceptable model fits according to CFI, but not the other indices (SRMR, *X*^2^, RMSEA). However, no standardized path effects were significant (Table 4).

## Discussion

The current study assessed the unique and joint contributions of hippocampal microstructure (diffusion) and macrostructure (volume) to age-related differences in multiple forms of episodic memory using SEM in a large sample of younger and older adults. In addition to replicating previously observed negative effects of age on hippocampal diffusion (Franco et al., 2021; Metzler-Baddeley et al., 2019; Nazeri et al., 2015; Radhakrishnan et al., 2020; Venkatesh et al., 2020), hippocampal volume (Du et al., 2006; Raz et al., 2005; Walhovd et al., 2005), and memory performance (Craik & McDowd, 1987; Nyberg, Bäckman, et al., 1996; Rönnlund et al., 2005), our results revealed two key findings. First, a significant indirect effect for hippocampal diffusion, but not volume, revealed that microstructure mediated the effect of age on memory performance, extending the literature by replicating similar findings from unimodal MRI studies within the same sample and using multi-compartment diffusion metrics. Second, the influential indirect pathway was not significant, indicating that hippocampal diffusion and volume made independent contributions to the age-episodic memory relationship.

Indirect effects revealed that the age-related decline in episodic memory was significantly mediated by hippocampal diffusion, but not hippocampal volume. These finding are consistent with previous unimodal multi-compartment diffusion-weighted (Radhakrishnan et al., 2020; Venkatesh et al., 2021) and T1-weighted (Raz et al., 1998; Rodrigue & Raz, 2004; Tisserand et al., 2000) studies that included younger and older adults. However, an advantage of the current multimodal approach is that these unique effects for each structural MRI modality were assessed within the same sample, revealing that diffusion was more sensitive to memory deficits in aging than volume. At least one multimodal imaging study within older adults similarly found that hippocampal diffusion, but not hippocampal volume, predicted episodic memory performance when using separate regression models for each structural modality (den Heijer et al., 2012), although they did not assess whether either structural metric mediated the effect of age on episodic memory, which was of interest here. The current study further extends these literatures by having used a latent memory construct that captured shared variance across multiple episodic memory metrics, not all of which were included in previous studies.

The non-significant influential indirect effect revealed that hippocampal diffusion and volume did not jointly contribute to the age-related decline in episodic memory. This finding is consistent with some multimodal MRI studies that focused on structure-memory relationships independent of age, which reported independent contributions of hippocampal diffusion and volume to episodic memory performance (den Heijer et al., 2012), but not with others whose approaches focused on the additive (Radhakrishnan et al., 2022) or shared (Köhncke et al., 2021) variance across these structural MRI measures. Similar results have also been observed in multimodal MRI studies that examined the contribution of volume in hippocampal gray matter and diffusion in white matter emanating from the hippocampus (e.g., fornix) to memory performance in aging (Foster et al., 2019; Gorbach et al., 2017; Hayek et al., 2020), with one study in adults across the lifespan finding that the effect of age on associative memory was mediated by fornix diffusion, but not volume of medial temporal structures that included the hippocampus (Foster et al., 2019). Finding similar relationships to age, but different relationships to episodic memory performance, supports the notion that diffusion– and T1-weighted MRI modalities are capturing at least some distinct neurobiological substrates in hippocampal gray matter (Wolf et al., 2015), or perhaps the same substrates but to different degrees. We speculate that microstructural metrics may be more influenced by subtle neuronal (e.g., neuron density, myelination, arborization) or glial (e.g., swelling) tissue properties than macrostructural metrics, making them more sensitive to individual– and age-related differences in memory performance.

One caveat of our study is that the small and non-significant contribution of hippocampal volume to the age-memory relationship across all participants may have affected our ability to detect an influential indirect effect. Unimodal T1-weighted studies in older adults have observed significant relationships between hippocampal volume and episodic memory (Bender et al., 2013; Bennett et al., 2019; Bruno et al., 2016; de Leon et al., 1997; Golomb et al., 1994; Lye et al., 2004; Shing et al., 2011), but the current sample of older adults may be too small to reliably test with our SEM model, as seen by there being no significant standardized path effects in the exploratory model within older adults only.

In summary, the current study revealed that, although both hippocampal macrostructure (volume) and microstructure (diffusion) are worse with age, only hippocampal microstructure contributed to worse episodic memory performance in older than younger adults. Our findings for NODDI diffusion metrics have implications for biomarker research aimed at establishing reliable neural metrics that vary with age and memory performance. We further demonstrate that SEM is an important tool for multimodal MRI, enabling the characterization of unique (specific indirect effects) and joint (influential indirect effect) contributions of each modality to cognitive aging.

## Acknowledgements

The authors wish to thank Justino Flores for his role in data collection. This work was supported by R00 AG047334 (Bennett) and R21 AG054804 (Bennett) from the National Institutes of Health/National Institute on Aging.

